# Random, fragile, or correlated: Mechanisms of synteny decay in mammals

**DOI:** 10.1101/2025.10.13.682123

**Authors:** Alexander S. Moffett, Michele Di Pierro

## Abstract

We formalize three models of genome evolution, yielding analytical distributions of synteny block sizes between pairs of genomes. To evaluate each model, we fit theoretical synteny block size distributions to empirical distributions from a comparative analysis of nearly 200 mammalian genomes. Between the random breakage model (RBM), with uniformly distributed breakpoints, the fragile breakage model (FBM) with breakpoint hotspots, and what we term the correlated breakage model (CBM) with pairs of nearby breakpoints, we find that the FBM and CBM are far more consistent with mammalian genome evolution than the RBM. Inferred parameters of each model qualitatively agree with previous characterizations of mammalian genome evolution. The FBM and CBM perform comparably with each other, suggesting that either model could serve as an improved null model of genome evolution to use in statistical tests for syntenic regions that are unlikely to exist due to random chance.

## 1. Introduction

Assisted by the abundance of chromosome-length genome assemblies now available, we have begun to understand how large-scale structural mutations shape the evolution of genomes. Much as conservation of a nucleotide or amino acid can be used to understand the evolution of genes or non-coding loci, the conservation of larger genomic regions, called microsynteny, provides a basis for studying genome evolution. In particular, the distribution of synteny block sizes observed when comparing two genomes can be used to extract information about the evolutionary pathway connecting the two genomes. Using genetic linkage maps, Nadeau and Taylor [1] developed a model (the “random breakage model”, or RBM) based on the assumption that disruptions between neighboring ancestral genes are uniformly distributed across the genome. They predicted that the lengths of synteny blocks, measured in centimorgans, would be exponentially distributed. Later work suggested that there are hotspots in genomes where evolutionary breakpoints are more likely to occur [2, 3, 4, 5, 6, 7] (the “fragile breakage model”, or FBM), providing evidence against uniformly distributed breakpoints. This heterogeneity of breakpoint rates is likely due to a combination of sequence features, structural features, and selection [5]. The FBM predicts an excess of short synteny blocks as compared with the exponential synteny block size distribution of the RBM, caused by the clustering of breakpoints in hotspots.

Neither the RBM nor the FBM account for clustering of breakpoints due to inversions, where the two breakpoints introduced by an inversion tend to be close to one another [8, 9, 10, 11, 12, 13, 14, 15]. While inversion-like double breakpoints are an integral part of analyses inferring the minimal number of rearrangements needed to transform one genome into another [16, 17, 2], realistic inversion length distributions are not taken into consideration in these studies. Similarly, several statistical models have been proposed including correlated breakpoints. Working with mammalian genomes, Berthelot *et al*. [18] developed and simulated a model where pairs of breakpoints occur with distances between them determined by a power law contact probability distribution from Hi-C experiments [19]. Restricting breakpoints to open chromatin regions, Berthelot *et al*. recovered a synteny block size distribution with an excess of short synteny blocks as compared with the random breakage model. Similarly, Biller *et al*. proposed a model with double breakpoints reflecting inversions [20], although they did not include any constraints on the inversion size distribution.

We refer to models with explicit consideration of inversions and their length distribution as examples of a correlated breakage model (CBM). The distinction between the FBM and the CBM lies in how regions with multiple nearby breakpoints arise. In the FBM, breakpoints are clustered due to their higher propensity in certain genomic regions. In the CBM, breakpoint clustering is a consequence of multiple breakpoints simultaneously occurring in close physical proximity, but not necessarily because they are in hotspots.

Despite this large body of work on models of genome evolution, often focused on reconstructing chromosomal rearrangement events and inferring ancestral gene orders [17, 21, 22, 23, 24, 25], analytical theory of genome evolution has lagged behind. The jump model describing prokaryotic genome evolution [26, 27, 28, 29] and models describing the statistics of gene co-localization on the same chromosome [30] have developed statistical theory beyond the RBM, but analytical work on the statistics of the CBM and FBM remains underdeveloped.

Here, we formulate analytical models to describe synteny block size distributions of the RBM, CBM, and FBM and fit them to a comparative analysis of 193 chromosome-length mammalian genome assemblies. Our goal is to evaluate the RBM, CBM, and FBM as null models of genome evolution which can be used to test specific hypotheses about the origins of syntenic regions. We provide simple expressions for the synteny block size distributions of each model while comparing the models on an equal footing from a statistical perspective. We confirm that breakpoint clustering, either due to inversion-like processes or hotspots, is crucial to accurately describing mammalian genome evolution. While the FBM performs better than the RBM or CBM, it requires an additional parameter over the CBM while providing a marginal improvement in goodness of fit. Our results suggest that the CBM serves as an improved neutral model over the RBM, while describing synteny block size distributions nearly as well as the FBM.

## 2 Results

Microsynteny is a property of a pair of genomes, which we define as regions of conserved gene order across the two genomes. If a set of genes is in the same order in the genomes of two individuals from species *A* and *B* (i.e. a synteny block), we can infer that those genes were in the same order in the last common ancestor of *A* and *B*, say *C*. Any rearrangement in either the lineage leading from *C* to *A* or *C* to *B* has the effect of breaking up synteny blocks into a greater number of smaller synteny blocks. In other words, synteny blocks between *A* and *B* are regions within which no breakpoints have occurred in either lineage while at least one breakpoint has occurred in either lineage at both ends of the region.

We model this process by treating the genome of *C* as an infinitely long linear chromosome which can be broken by chromosomal rearrangements that occur according to a Poisson process. The size distribution of synteny blocks is determined by the distances between neighboring breakpoints (measured in number of genes), so we can recover these distributions by conditioning on a breakpoint having occurred in a single inter-gene region and calculating the probability that exactly *n* − 1 neighboring inter-gene regions have had zero breakpoints occur within them in the total evolutionary time traversed in the independent lineages *C* to *A* and *C* to *B, t* = *t*_*CA*_ + *t*_*CB*_. We use the mean amino acid substitutions per site (abbreviated as subs/site) across homologs, as calculated by OrthoFinder [31], as a rescaled unit of evolutionary time.

We use the term “cut” as a stand-in for “breakpoint” when describing our mathematical models. A 1-cut is simply a single breakpoint that forms independently of any other breakpoint. If there is an equal probability for a 1-cut to occur within any inter-gene region, we have a version of the RBM, which we call the 1-cut model here (Fig. 1a). In this model, 1-cuts form at a rate *κ* in units of sites*/*(sub *×* inter-gene region), so that the probability of at least one 1-cut occurring in a given inter-gene region in the total evolutionary time elapsed between *A* and *B* is

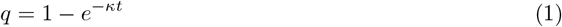

found from a Poisson distribution with parameter *κt*. We can then use Eq. 1 as a parameter in a geometric distribution to find the probability that exactly *n* genes will remain in microsynteny (or equivalently, that exactly *n* − 1 inter-gene regions will have zero cuts)

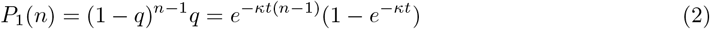

**Figure 1:**
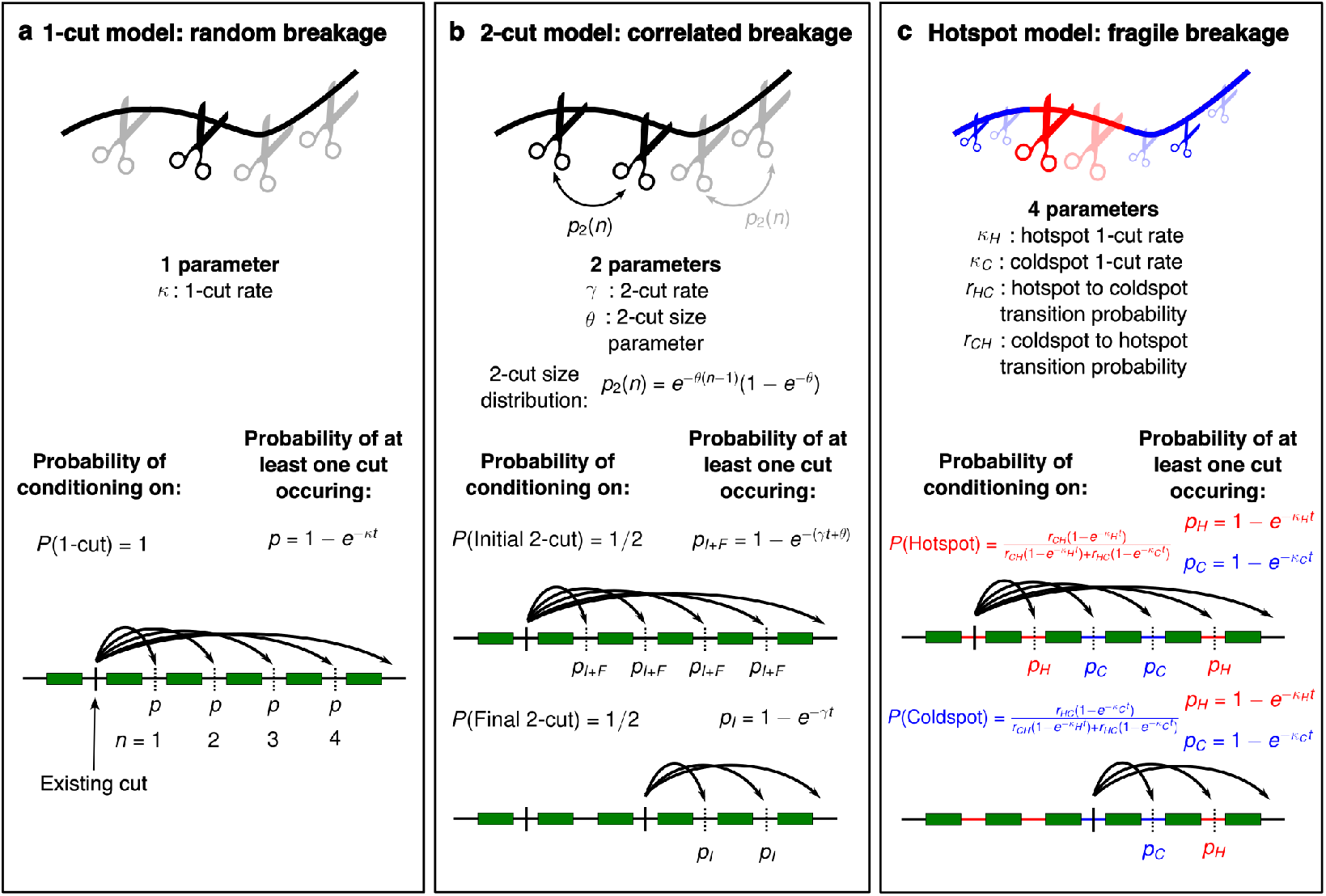
Models of synteny in genome evolution. The 1-cut model (**a**) corresponds to the random breakage model. We call a single breakpoint a 1-cut, where the probability of a 1-cut occurring at any single inter-gene region is constant. For two species separated by *t* average substitutions per site, the probability of at least 1-cut occurring in any given inter-gene region is equal to 1 − *e*^−*κt*^, where *κ* is the 1-cut rate. By assuming that at least one 1-cut has occurred (the reference cut), we can calculate the probability that a region of *n* neighboring genes will remain in synteny, uninterrupted by breakpoints. Since there is only one type of breakpoint in this model, the probability of conditioning on a 1-cut is *P* (1-cut) = 1. In the 2-cut model (**b**), there are now two types of breakpoints. A 2-cut occurs when two breakpoints occur simultaneously on the same chromosome, for example inversions spanning multiple genes. Similarly to a 1-cut, the probability of the initial breakpoint of a 2-cut (initial 2-cut) forming at any inter-gene region is 1 − *e*^−*γt*^, where *γ* is the 2-cut rate. A second breakpoint (final 2-cut) then occurs according to a geometric distribution (*p*_2_ (*n*)) with probability of breakage at each inter-gene region 1 − *e*^−*θ*^ . Each type of 2-cut makes up half of all breakpoints, so initial and final 2-cuts are conditioned on with probability 1*/*2 each, which can be written as *P* (Initial 2-cut) = *P* (Final 2-cut) = 1*/*2. The hotspot 1-cut model (**c**) consists of two types of 1-cuts, in hotspots and coldspots. Hotspots and coldspots are modeled as a Markov chain with transition probability from hot to cold adjacent inter-gene regions *r*_*HC*_ and cold to hot regions *r*_*CH*_ . Breakpoints occur in hotspots at rate *κ*_*H*_ and in coldspots at rate *κ*_*C*_, where *κ*_*H*_ ≥ *κ*_*C*_ . Conditioning on a hotspot occurs with probability *P* (Hotspot) = *r*_*CH*_ *κ*_*H*_ */*(*r*_*CH*_ *κ*_*H*_ +*r*_*HC*_ *κ*_*C*_) and on a coldspot with probability *P* (Coldspot) = *r*_*HC*_ *κ*_*C*_ */*(*r*_*CH*_ *κ*_*H*_ + *r*_*HC*_ *κ*_*C*_).

This geometric distribution is the discrete equivalent of the exponential RBM distribution derived by Nadeau and Taylor [1]. Note that the 1-cut model requires one parameter.

To formulate a statistical model of the CBM, we need to introduce a new type of cut, the 2-cut (Fig. 1b). When a 2-cut occurs, two cuts necessarily form on the same chromosome with the number of genes between them sampled from a 2-cut size distribution. We formulate 2-cuts as occurring in two steps. First an initial 2-cut occurs just as like a 1-cut, at an equal probability for each intergene region according to rate *γ*. Then, a final 2-cut is chosen *n* genes away from the initial 2-cut, according to a geometric distribution with parameter 1 − *e*^−*θ*^. Because there are two subtypes of 2-cut, when we condition on an existing 2-cut we may be conditioning on either an initial or final 2-cut, each with probability 1*/*2 because they occur in equal numbers. If we have conditioned on an initial 2-cut, either the associated final 2-cut or another independent initial 2-cut can disrupt microsynteny. If instead we have conditioned on a final 2-cut, only an independent inital 2-cut can disrupt local microsynteny, neglecting the possibility of certain overlapping 2-cut regions (see Methods for more detail). This leads to the 2-cut synteny block size distribution

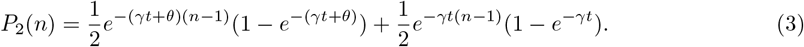

The 2-cut model requires two parameters, while the 1-cut and 2-cut models can be combined into a three parameter model allowing for both cut types (see Methods).

We describe the FBM with a modified, four-parameter version of the 1-cut model, which we call the hotspot model (Fig. 1c). We consider two types of inter-gene regions, either hotspots or coldspots, where 1-cuts occur with rates *κ*_*H*_ and *κ*_*C*_ with *κ*_*H*_ ≥ *κ*_*C*_. We assume the distribution of hotspots and coldspots along chromosomes can be described by a Markov chain with probabilities *r*_*HC*_ and *r*_*CH*_ of transitioning from a hot to a coldspot, and vice versa, between neighboring inter-gene regions. The probabilities of conditioning on a hotspot or coldspot are determined by all four parameters *κ*_*H*_, *κ*_*C*_, *r*_*HC*_, and *r*_*CH*_ . See the Methods section for the hotspot synteny block size distribution, as well as more detail on the other models.

In order to validate the theoretical distributions of each model, we ran 100 independent stochastic simulations each of the 1-cut, 2-cut, and hotspot models. We started each simulation from a synthetic, ancestral genome with 20 chromosome each containing 1000 genes, and ran them for 0.25 subs/site using realistic but arbitrarily chosen parameters. Because the theoretical distributions do not use any information about the ancestral genome, at short times the theoretical distributions for each model predict a lower probability of long synteny blocks containing 1000 genes than is found in simulations (Fig. 2). Even so, the theoretical models qualitatively match the shape of the synteny block size distributions for shorter blocks. At longer times, the theoretical and simulation-derived distributions closely match.

**Figure 2:**
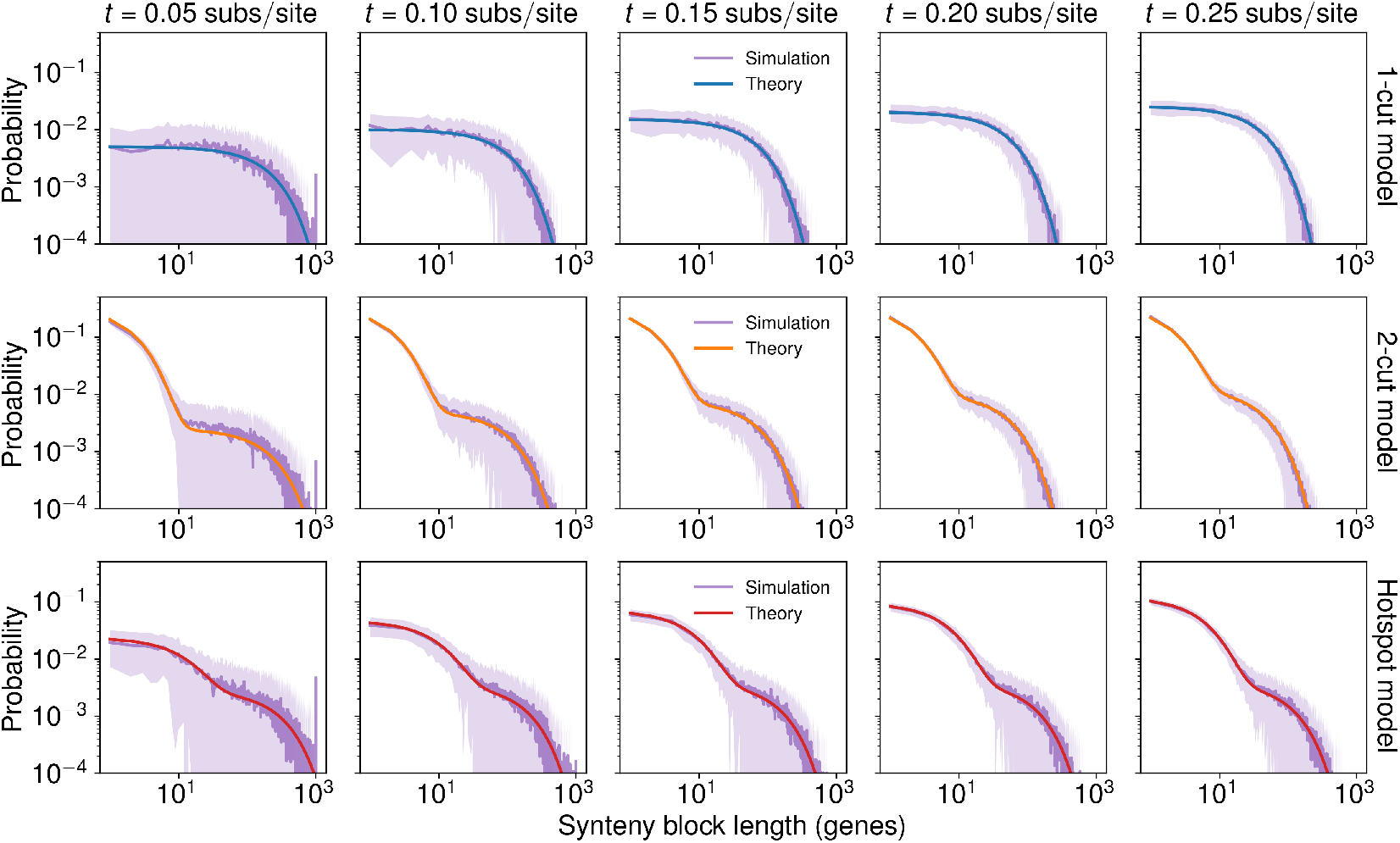
Analytical theory describes long-term synteny block size distributions. Comparison of synteny block size distributions between stochastic simulations of genome evolution with analytical models for the 1-cut model (**first row**), 2-cut model (**second row**), and hotspot model (**third row**). The initial genome for each simulation consists of 20 chromosomes each with 1000 genes. We ran 100 independent simulations of each model. Dark purple lines are the mean probability, while the lighter purple shaded areas show the standard deviation in probability. At short times (0.05 subs/site for all three models and 0.1 subs/site for the hotspot model) there is a peak in the simulations at synteny blocks with 1000 genes, reflecting initial conditions. The analytical theory by design lacks information about initial conditions and matches relatively poorly with simulations at these short times as compared with simulations at later times. Even with the disagreements between theory and simulations at short times and long synteny blocks, theoretical distributions qualitatively match with distributions from simulations. Once the probability of large synteny blocks is very low (i.e. the influence of initial conditions is low), the analytical models match well with simulations. We used *κ* = 0.1 sites/sub for the 1-cut model, *γ* = 0.1 sites/sub and *θ* = 0.5 for the 2-cut model, and *κ*_*H*_ = 0.5 subs/site, *κ*_*C*_ = 0.01 subs/site, *r*_*HC*_ = 0.05, and *r*_*CH*_ = 0.01 for the hotspot model.

Next, we fit each model, including the 1-cut + 2-cut model, to our comparative analysis of mammalian genomes. We used the microsynteny analysis from Moffett *et al*. [32] of 191 mammalian genome assemblies and a single bird outgroup genome assembly from the DNA Zoo [33] and human and mouse genomes from GENCODE [34]. Using the APES algorithm [32] to determine microsynteny between pairs of genomes, we determined the distribution of synteny block sizes, measured by the number of genes in a synteny block, for each genome pair. To fit each model, we numerically minimized the Kullback-Liebler divergence between theoretical and empirical synteny block size distributions inde-pendently for each genome pair. The four models we discuss in this article are nested models with one (1-cut), two (2-cut), three (1-cut + 2-cut), and four (hotspot) parameters (Fig. 1). For this reason, we can guarantee that each model should perform at least as well as the model with one fewer parameters.

The geometric distribution predicted by the 1-cut model underestimates the number of short synteny blocks, while the 2-cut, 1-cut + 2-cut, and hotspots models can each qualitatively match the distribution of short synteny blocks accurately (Fig. 3a-c). Considering all genome pairs, the 1-cut distribution consistently performed the worst of all models, with much larger Kullback-Leibler divergences (Fig. 3d). The 2-cut and 1-cut + 2-cut models performed almost identically, indicating that little is gained by allowing for both 1-cuts and 2-cuts (Fig. 3d). The hotspot model tended to achieve the lowest Kullback-Liebler divergences, although there is significant overlap between the distributions of hotspot and 2-cut Kullback Liebler divergences (Fig. 3d). While using alternate APES parameters that tend to either increase or decrease the length of synteny blocks noticeably alter the empirical and theoretical synteny block size distributions, the relative performances of the four models remained qualitatively unchanged (Figs. S1 & S2).

**Figure 3:**
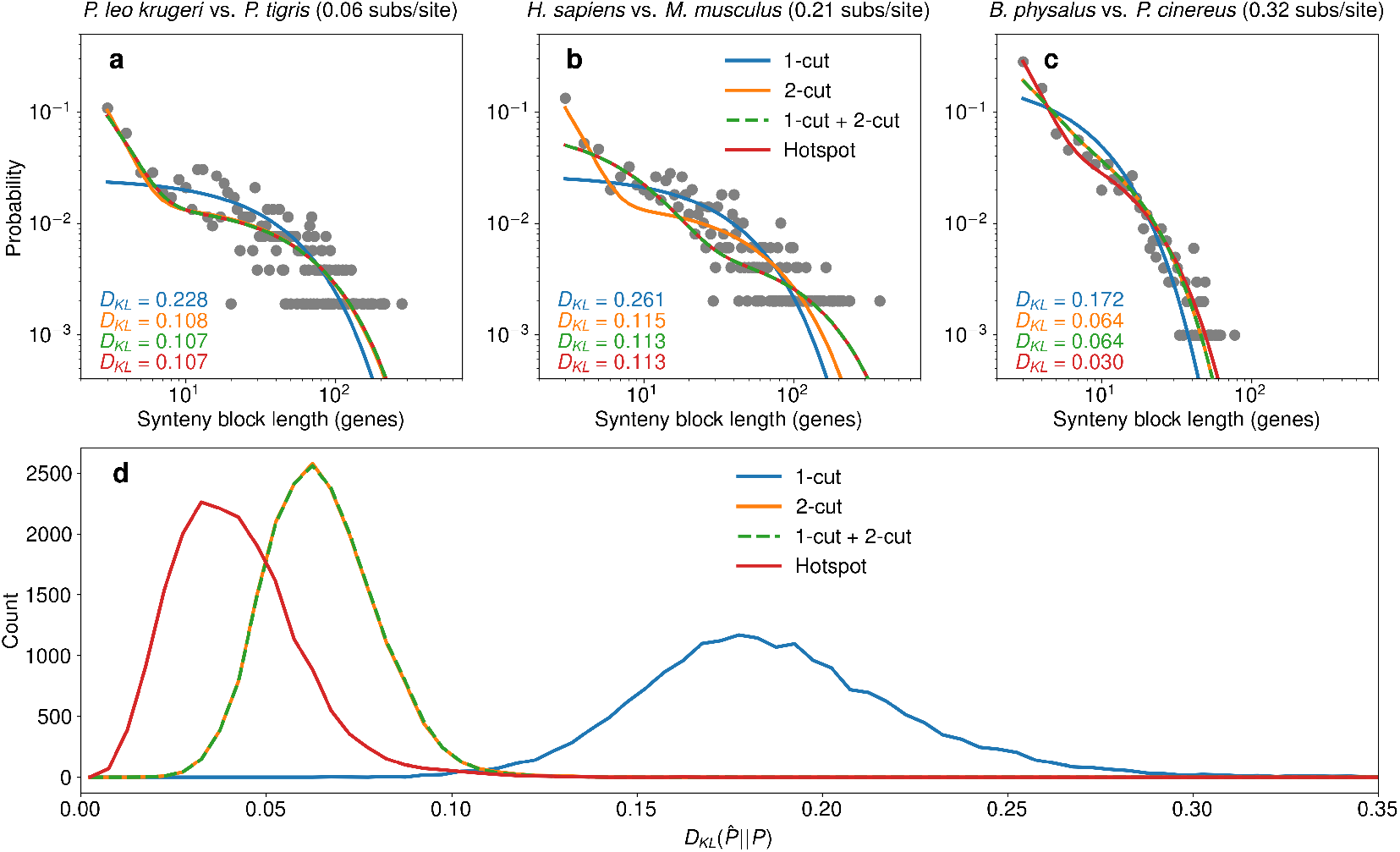
Comparing fits of theoretical models to data. Synteny block length distributions (gray dots) calculated from comparison between (**a**) *P. leo krugeri* vs. *P. tigris*, separated by 0.06 amino acid substitutions per site on average, (**b**) *H. sapiens* vs. *M. musculus*, separated by 0.21 amino acid substitutions per site on average, and (**c**) *B. physalus* vs. *P. cinereus*, separated by 0.32 amino acid substitutions per site on average. For each comparison, we show fits of theoretical distributions from the 1-cut model, 2-cut model, 1-cut + 2-cut model, and hotspot model. Color-coded Kullback-Liebler divergences (*D*_*KL*_) between each fit distribution and the empirical distribution are shown in the bottom lefthand corner of each subplot. The hotspot 1-cut model performs among the best models in all three cases, while the neutral 1-cut model cannot account for the observed excess of short synteny blocks. (**d**) Distributions of *D*_*KL*_ between model fits and empirical synteny block length distributions across all 18,721 unique species pairs. The neutral 1-cut model has much larger *D*_*KL*_ values than any other model, while the neutral 2-cut and neutral 1-cut + 2-cut model achieve essentially the same *D*_*KL*_ values. The hotspot 1-cut model *D*_*KL*_ distribution is shifted slightly to lower values than the neutral 2-cut and neutral 1-cut + 2-cut models, indicating moderate improvement by the hotspot 1-cut model. The APES parameters used for this figure are 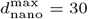 and 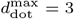.

The inferred parameters for the 1-cut, 2-cut, and hotspot model make specific predictions about the distributions and rates of breakpoints in mammalian evolution. Fits of the 1-cut model yield a sharply peaked distribution of 1-cut rates close to 0.5 sites/sub (Fig. 4a). The inferred 2-cut rates are bimodal, with most rates being very small while a smaller number of inferred rates are peaked around 0.3 sites/sub (Fig. 4b). The bimodality of the *γ* distribution is likely due to the fact that the 2-cut model begins to act more like the 1-cut model with small *γ* values. Given the separation of Kullback-Liebler divergences between the 1-cut and 2-cut models (Fig. 3d), it is clear that the 2-cut model is almost always behaving differently from the 1-cut model. Still, the small *γ* values are likely due to cases where the synteny block size distribution is in some sense reflecting a “1-cut-like” process. The distribution of the 2-cut size parameter *θ* is also tightly peaked, corresponding to a mean 2-cut size of 2.62 genes or 236 kb (see Fig. S3 for details of estimating 2-cut size in base pairs). The inferred hotspot model parameters reflect very low 1-cut rates in coldspots, with the vast majority of *κ*_*C*_ values falling close to zero, although a small fraction of genome pairs have a larger *κ*_*C*_ value close to 0.25 sites/sub (Fig. 4c). The hotspot cut rate *κ*_*H*_, has a peaked distribution around 3 sites/sub, corresponding to an average distance between hotspot 1cuts of 2.05 genes or 185 kb (Fig. 4c). This mean inter-cut distance is comparable with that predicted by the 2-cut model, suggesting that 1-cuts within hotspots and 2-cuts capture similar aspects of the synteny block size distribution. For most genome pairs, the majority of inter-gene regions are predicted to be coldspots (Fig. 4d), in agreement with past estimates [35]. We also find that the mean hotspot length, defined as the mean number of consecutive hotspot inter-gene regions (1*/r*_*HC*_) times a constant conversion factor of 90 genes/kb (Fig. S3), falls below 1 Mb for almost all genome pairs (Fig. 4e). This is in agreement with previous predictions of the FBM [7]. We predict coldspots to be larger, typically falling between 1 Mb and 2 Mb (Fig. 4e).

**Figure 4:**
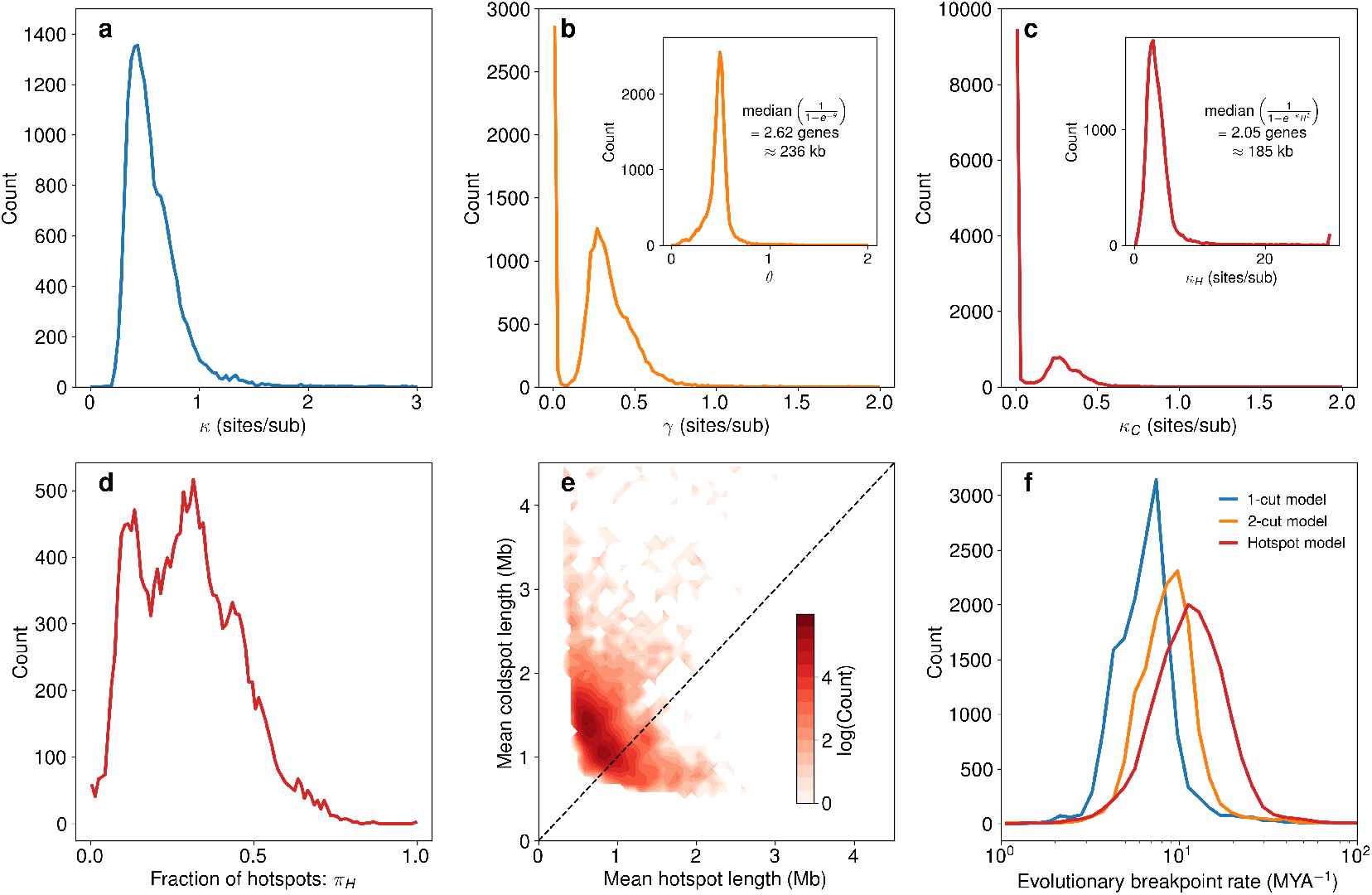
Inferred model parameters for mammals. We show distributions over all 18,721 pairwise genome comparisons for each parameter inferred for each model. For the 1-cut model (**a**), the vast majority of inferred 1-cut rate (*κ*) values fell between 0 and 1 sites/sub. For the 2-cut model (**b**), 2-cut rates (*γ*) are either near zero or are peaked around 0.3 sites/sub. The inset subfigure shows the distribution of inferred *θ* values, which are tightly peaked close to 0.5, with a median 2-cut size of 2.62 genes or approximately 236 kb (see Fig. S5 and SI Methods for details on conversion between genes and base pairs). Inferred coldspot 1-cut rates (*κ*_*C*_) are typically close to zero (**c**), representing regions with very few breakpoints, while in a smaller subset of genome comparisons coldspot 1-cut rates peaked around 0.25 sites/sub. In the inset subfigure, the distribution of hotspot 1-cut rates (*κ*_*H*_) is tightly peaked around 3 sites/sub, with a mean distance between 1-cuts within hotspots of 2.05 genes or approximately 185 kb. The inferred fraction of inter-gene regions that are hotspots (**d**) is less than 0.5 for most genome comparisons, meaning that most inter-gene regions are coldspots. Analysis of the lengths of hotspots and coldspots in each genome comparison (**e**) shows that hotspots are typically much smaller than coldspots, where most hotspots are smaller than 1 Mb and coldspots fall between 1-2 Mb. Distributions of mean evolutionary breakpoint rates (**f**) show increasing rates between the 1-cut, 2-cut, and hotspots models, with most rates falling between 1 MYA^−1^ and 50 MYA^−1^ (see SI Methods for details). The APES parameters used for this figure are 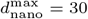 and 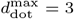.

Inferred evolutionary breakpoint rates, the average number of breakpoints predicted from model parameters for a pair of genomes divided by twice the time to their last common ancestor (see Methods), increased from the 1-cut model to the 2-cut model and from the 2-cut model to the hotspot model (Fig. 4f). The predicted values fell between about 2 MYA^−1^ to 70 MYA^−1^, exceeding reported breakpoint rates in mammals by at most two orders of magnitude [24]. We note that the breakpoint rates predicted here are for the whole pathway between species in the phylogenetic tree, while Damas *et al*. report breakpoint rates by branch [24].

While *κ, γ*, and *κ*_*C*_ all shift to larger values as APES parameters are shifted to values promoting larger numbers of shorter synteny blocks, *θ* and *κ*_*H*_ are intriguingly insensitive to APES parameter choices (Fig. S4). This suggests that the small rearrangements found in our analysis cannot easily be dismissed as artifacts of the APES algorithm. On the other hand, the fraction of inter-gene regions that are hotspots depends strongly on APES parameter choices (Fig. S5). For APES parameters that favor fewer, long synteny blocks, the predicted breakpoint rates are much closer to those inferred by [24] (Fig. S5).

## 3 Discussion

We have derived simple, analytical models for the synteny block length distributions resulting from four models of genome evolution and tested them against a large comparative analysis of mammalian genomes. We confirm arguments that the RBM (1-cut model) is inadequate for describing mammalian genome evolution, while providing extensive evidence that the CBM (2-cut model) and FBM (hotspot model) perform far better, but comparably with each other. We find that an excess of small synteny blocks (∼ 3−10 genes) over predictions of the 1-cut model, in agreement with previous work [36, 7, 18], is likely universal across pairwise comparisons of mammalian genomes suggesting a common mechanism throughout mammal evolution. Both the 2-cut and hotspot models succeed in describing this phenomenon, meaning that clustered breakpoints, as one might expect with inversions, concentration of single breakpoints in hotspots, or some mixture of the two, could be responsible. Independent evidence for inversion size distributions favoring smaller inversions [8, 9, 10, 12, 15] and breakpoint hotspots [2] support the idea that either mechanism could plausibly contribute to synteny block length distributions.

All four models we have tested are coarse descriptions of genome evolution, reflecting the statistical consequences of genome rearrangements rather than the complicated details of the evolutionary histories of specific chromosomal regions. Our goal was to evaluate these coarse-grained theories to then choose the best null model of genome evolution against which more detailed hypotheses can be tested. Our analysis suggests that either of the 2-cut or hotspot models can be used in this capacity, as both include mechanisms for generating the excess of short synteny blocks as compared with the 1-cut model. The combined neutral 1-cut + 2-cut model appears to have few advantages over the 2-cut model.

Crucially, the hotspot model is not a neutral model, in the sense that it allows for variation of 1-cut rates along chromosomes. The current analysis says nothing conclusive about whether selection or other processes are present or absent, only that from the synteny block size distributions, variations between hotspots and coldspots looks very similar to inversion-like correlated breakpoints. Moreover, by design our analysis says nothing about specific locations of hotspots and coldspots on ancestral genomes, but rather predicts plausible statistical properties of hotspots and coldspot distributions across genomes.

To simplify our analysis, we have assumed that breakpoints are always in inter-gene regions and while we ignore the length of inter-gene regions. Lemaitre *et al*. found evidence that most breakpoints are in inter-gene regions [37], in line with predictions that purifying selection should often prevent breakpoints within protein coding genes. Biller *et al*. [20] pointed out possible issues with ignoring inter-gene size, which they call the pseudouniform breakage model in contrast to a true uniform breakage model where a neutral model would assign breakage rates proportional to inter-gene region lengths. It is unclear if the uniform breakage model of Biller *et al*. actually corresponds to the original model of Nadeau and Taylor [1], who measured distances between genes from linkage maps in units of centimorgans. Heterogeneity in recombination rates would disrupt proportionality between genetic linkage map distances and genomic distances measured in basepairs. Further, Berthelot *et al*. [18] showed that the average number of breakpoints within inter-gene regions weakly depends on the size of the region, according to 2.4 *·* 10^−3^ *· L*^0.28^ where *L* is the inter-gene region size in basepairs. This is evidence against proportionality between breakpoint rate within an inter-gene region and the size of the region, suggesting that ignoring inter-gene region sizes is a reasonable approximation.

Repetitive sequences and segmental duplications [3, 38, 39], 3D genome structure [18, 40, 41, 42, 43], and selection [37, 35] all likely play a role in determining the patterns of breakpoints in mammals [44]. We have deliberately left out these causal factors from our models, which are meant to serve as null models of genome evolution. Our formulations of the CBM and the FBM can be adapted to test for deviations from expected behavior in order to identify statistically significant associations between orthogroups across mammalian evolution. Our models can also be extended to account for phylogeny, allowing for inference of model parameters on each branch of a species tree. While the analysis presented here is on pairs of genomes which are not phylogenetically independent, the separation of Kullback-Leibler divergence distributions between the different models presents clear support for the CBM and FBM models. Branch dependent extensions of our models will remove dependence between pairwise comparisons and provide further insight into evolutionary histories.

## 4 Methods

### 4.1 Genome data and orthogroup detection

We used MAKER2 [45] genome annotations of 191 mammals and 1 bird (as an outgroup) from the DNA Zoo [33] in addition to GENCODE human genome annotation v43 and mouse genome annotation v32 [34]. As described in our previous work [32], we used OrthoFinder 2 [31, 46] to determine orthogroups and to infer a rooted species tree [47, 48].

### 4.2 Microsynteny inference with APES

We used the APES algorithm [32] to infer microsynteny between all pairs of genomes in our dataset from dot plots. APES first finds regions of perfectly conserved gene order (nanosynteny) and then connects statistically significant nanosynteny blocks into microsynteny blocks. We use a minimum nanosynteny block size of three genes. Two parameters control how nanosynteny blocks are connected: 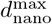, which sets a limit on how large gaps between nanosynteny blocks can be from one another, and 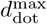, which controls how large gaps between dots can be within microsynteny. Increasing 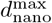 and 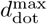 means that more nanosynteny blocks will be connected into larger microsynteny blocks. Note that here the size of a synteny block refers to the number of gene pairs in that block. See [32] for a detailed description of APES. Python code is available at https://github.com/DiPierroLab/GenomicTools.

### 4.3 Synteny block size distribution models

We derive synteny block size distributions from one species, *A*, to another, *B*, by assuming that at least one breakpoint has formed along the path of lineages linking the two genomes. The length of this path is measured in average amino acids substitutions per site, *t*, which is related to the total time of independent evolution for the lineages of *A* and *B* since their most recent common ancestor. The distribution of synteny block sizes is then determined by the pattern of other breakpoints around this arbitrarily chosen reference breakpoint. Due to the assumption of uniform breakpoint probabilities across all inter-gene regions in random breakage and correlated breakage models, we are free to choose an arbitrary reference point and apply the results we derive to any point in an ancestral genome. For the fragile breakage model, where the breakpoint rate varies between hot and coldspots, we develop a mean-field theory that allows us to apply results from an arbitrarily-chosen reference breakpoint.

#### 4.3.1 1-cut model

The one parameter 1-cut model (Fig. 1b) corresponds to the random breakage model [1], except measuring distances in terms of numbers of genes instead of centimorgans. The number of 1-cuts occurring at any one inter-gene region on the species tree path between species *A* and *B, k*, is Poisson distributed according to

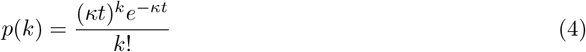

where *κ* is the rate of 1-cuts. Then, the probability of no cuts occurring in a particular inter-gene region on this path is

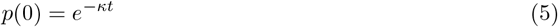

and the probability of at least one cut occurring is

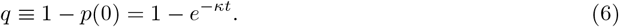

As mentioned above, the probability of observing a synteny block containing with *n* genes can be found by conditioning on a cut occurring at a reference inter-gene region, and then calculating the probability of no cuts occurring in *n*− 1 consecutive adjacent inter-gene regions followed by a cut occurring at the next inter-gene region

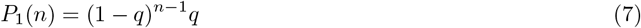

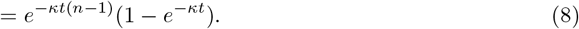

This is an approximation because we must assume that *t* is large enough so that the influence of the initial synteny block size distribution is negligible.

#### 4.3.2 2-cut model

A 2-cut involves in initial cut, chosen uniformly from all inter-gene regions, followed by a final cut with a geometrically distributed distance from the initial cut. Initial and final 2-cuts always come in pairs. This mimics inversions, though we remain agnostic as to the mechanism of breakpoint formation. Just like in the 1-cut model, one or more initial 2-cuts occur at a given inter-gene region according to

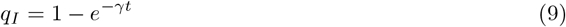

where *γ* is the 2-cut rate. Although this is the probability of one *or more* initial 2-cuts occurring at an inter-gene region, we assume that only one initial 2-cut occurs. We justify this approximation below.

Given that an initial 2-cut has occurred, a final 2-cut can then occur at nearby inter-gene regions with probability

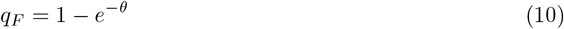

Note that this does not come from a Poisson distribution, we have simply defined *q*_*F*_ using the parameter *θ* to make *γ* and *θ* easily comparable. A final 2-cut will occur *n* genes away, or equivalently *n* − 1 inter-gene regions away, from its corresponding initial 2-cut with probability

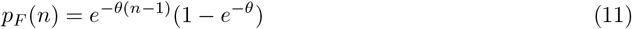

The choice of which step of a 2-cut is “initial” and which is “final” is arbitrary, and for convenience we adopt the convention that final 2-cuts always occur to the “right” of initial 2-cuts (when viewed as in Fig. 1).

Because initial and final 2-cuts occur in equal numbers, the probability that the inter-gene region we condition on is an initial 2-cut is equal to the probability that it is a final 2-cut, each 1*/*2. An initial 2-cut can have the closest cut be either its associated final 2-cut or another initial 2-cut, while a final 2-cut can only have another initial 2-cut as the closest cut. This implies an assumption about overlapping 2-cuts, that we will explore below. The 2-cut model distribution is then

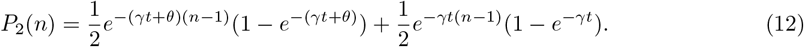

where the first term accounts for conditioning on an initial 2-cut and the second term for conditioning on a final 2-cut.

The region enclosed by one pair of initial and final 2-cuts can intersect with that enclosed by another pair of initial and final 2-cuts. A 2-cut with both the initial and final 2-cuts to the right of the reference cut occurs with probability (1 − *e*^−*γt*^) per inter-gene region. A 2-cut with an initial 2-cut on one side of the reference cut and a final 2-cut on the other side is more complicated to deal with. The probability that this occurs with a 2-cut a *n* genes away from the reference cut is

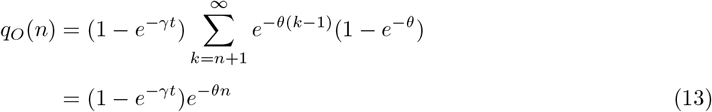

where (1 − *e*^−*γt*^) is the probability of an initial 2-cut forming, while the sum accounts for each corresponding final 2-cut that could cause a cut *n* genes away from the reference cut. We can combine Eq. 13 with Eqs. 9 & 10 to find an overall probability of a cut occurring to the right of an initial 2-cut

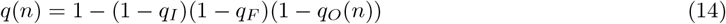

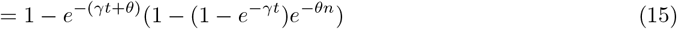

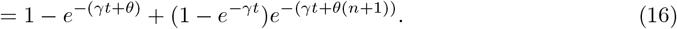

If *θ* is large enough, we can approximate *q*(*n*) as

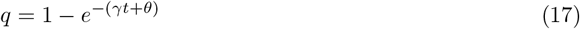

so that it no longer depends on *n*. An analogous derivation can be carried out for final 2-cut reference inter-gene regions. This holds when *q*_*O*_(*n*) is small enough, and because *q*_*O*_(*n*) is monotonically decreasing, we calculate

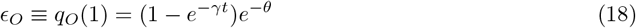

to assess the error in our model due to overlapping 2-cuts. As can be seen in Fig. S6a, this probability is typically low for models fit to our comparative analysis, with a median value of 0.034. This indicates that our models, as fit to the data, are largely internally consistent.

A second assumption we have made is that either zero or one initial 2-cuts occur at each inter-gene region. The probability of two or more initial 2-cuts occurring at any inter-gene region is

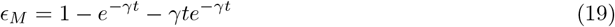

so that if *ϵ*_*M*_ is small, our assumption is reasonable. Again, with the inferred parameters in our analysis, *ϵ*_*M*_ is typically small, with a median value of 0.002, meaning our fits are internally consistent with respect to the assumption that no more than one initial 2-cut occurs at each inter-gene region (Fig. S6b).

#### 4.3.3 1-cut + 2-cut model

If we allow for both 1-cuts and 2-cuts to occur, we have a three parameter model very similar to the neutral 2-cut distribution, except that the probability of conditioning on an initial 2-cut is now 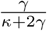 and the probability of conditioning on a 1-cut or final 2-cut is 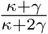 (Fig. 1d). This is because the average number of 1-cuts is *κt*, the average number of initial 2-cuts is *γt*, and the average number of final 2-cuts is *γt*. The synteny block size distribution is

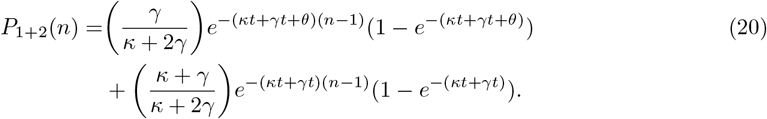

This reduces to the 2-cut model when *κ* = 0 and the 1-cut model when *γ* = *θ* = 0.

#### 4.3.4 Hotspot model

Finally, we consider a four parameter model where there are “hotspots” and “coldspots” for breakpoints, both occurring as 1-cuts (Fig. 1e). We model the distribution of hotspots and coldspots along a chromosome according to a two-state Markov chain with transition probability matrix

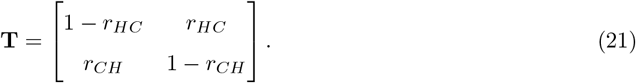

The parameters *r*_*HC*_ and *r*_*CH*_ are the probabilities of transitioning from a hotspot to a coldspot (coldspot to hotspot) when moving from one inter-gene region to an adjacent inter-gene region. The equilibrium probabilities of an inter-gene region being a hotspot or a coldspot are

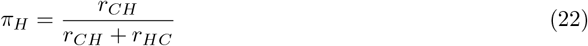

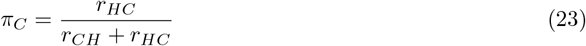

while the probability of a cut being in a hotspot

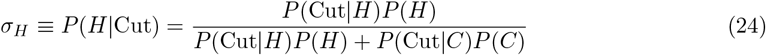

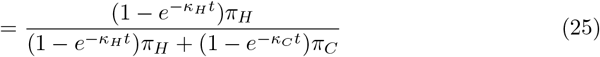

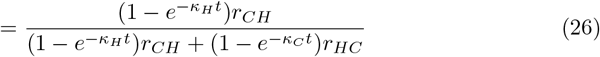

and in a coldspot

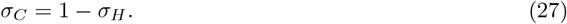

where the second equality approximately holds for small *κ*_*H*_*t* and *κ*_*C*_*t*. We write this approximate distribution as a row vector

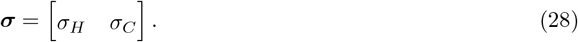

We also arrange the probabilities of no cut occurring in hotspots and coldspots in the matrix

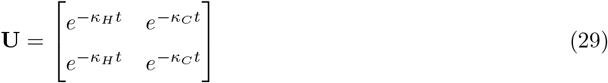

as well as the probabilities of at least one cut occurring in hotspots and coldspots

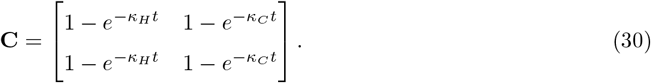

All together, we find the synteny block size distribution for the hotspot model to be

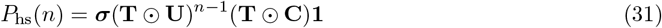

where ⊙ is the Hadamard product and **1** is a 2 *×* 1 column vector with both components equal to one.

### 4.4 Model fitting

For each pair of genomes, we use the Kullback-Liebler divergence between the empirical synteny block size distribution, 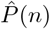, and the distribution predicted by a given model with parameter vector **x**, *P* (*n*; **x**), as an objective function for fitting parameters. The Kullback-Liebler divergence is defined as

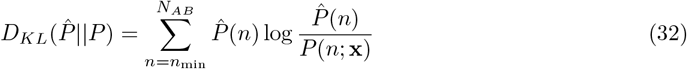

following the convention 0 *·* log 0 = 0. *P* (*n*) and 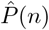 are normalized so that

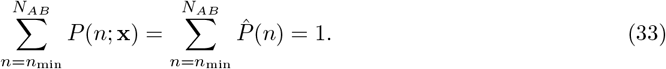

We use *n*_min_ = 3, as justified in our previous work [32]. We used the sequential least squares programming algorithm [49] implemented in Scipy [50] to minimize Kullback-Leibler divergences with respect to model parameters separately for each pair of genomes. In all cases, the gradient of the objective functions was calculated numerically. For each fit, we initiated optimization from 50 randomly chosen initial parameter vectors. Each parameter was chosen independently from all others from a uniform distribution over its allowed interval of values,

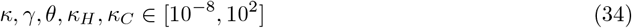

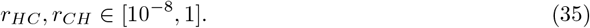

The parameters that achieved the smallest Kullback-Liebler divergence were taken to be optimal.

### 4.5 Simulations

All simulations were run on a synthetic genome with 20 chromosomes, each containing 1000 genes. We represent genomes as a graph with 20 connected components, each representing a chromosome. Each connected component has 1000 nodes, each representing a gene, where 998 nodes have degree two, while the two nodes on either end of the connected component have degree one. An edge between two nodes represents adjacency of two genes on a chromosome. We write this initial graph as

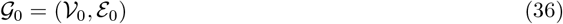

with |𝒱_0_| = 20, 000 and |ℰ_0_| = 19, 980 and with edges ℰ = *{*1, 2, 3, …, 19, 980*}*. In each simulation, we iterate between sampling a timestep, determining the breakpoints, and removing the affected edges from the graph. The evolving graph is written as

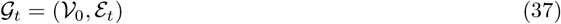

where the node set does not change with time. For the 1-cut model, timesteps are sampled according to

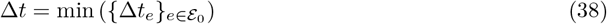

where the time to a cut in each edge is sampled independently from

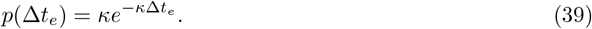

For the 2-cut model, this distribution becomes

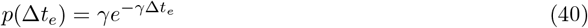

For the hotspot model, we first assign each edge in ℰ_0_ to be a hotspot or coldspot by sampling from the Markov chain represented by Eq. 21, with hotspot edges ℋ_0_ and coldspot edges *C*_0_ satisfying ℋ_0_ ∪ 𝒞_0_ = ℰ_0_ and ℋ_0_ ∩ 𝒞_0_ = ∅. Then, timesteps are chosen according to

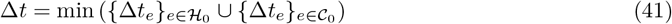

with

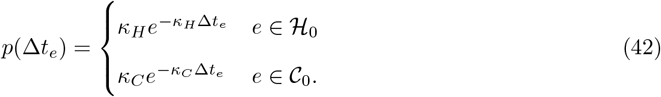

In the 1-cut and hotspot models, the edge *e* corresponding to the smallest timestep Δ*t*_*e*_ is removed from the edge set to yield

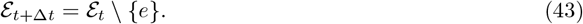

The 2-cut model requires a final 2-cut with each initial 2-cut, which is determined by sampling from the 2-cut size distribution

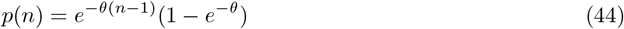

introducing a breakpoint *n* genes away from the initial 2-cut at edge *e*^*′*^ = *e* + *n* if *e* and *e*^*′*^ are in the same connected component in ℰ_0_. In this case, we update the edge set according to

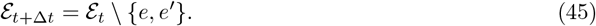

Otherwise if *e* and *e*^*′*^ are on different connected components in ℰ_0_, we update the edge set according to Eq. 43.

Starting from *t* = 0, time is updated to *t* ← *t* + Δ*t* at each timestep until *t* exceeds the maximum simulation time of 0.25 subs/site. At the end of each simulation, the distribution of connected component sizes in the graph at *t* = 0.25 subs/site is the distribution of synteny block sizes. We ran 100 independent simulations for each of the 1-cut, 2-cut, and hotspot models, and calculated the mean and standard deviation of the probabilities.

### 4.6 Inferring breakpoint rates

We used TimeTree [51] to find divergence times (*t*_*MY A*_) between species in millions of years. Not all species in our dataset are included in TimeTree, so missing species were excluded from this analysis. We use the total number of genes in synteny between two genomes *A* and *B* (*S*) as a estimate of the number of inter-gene regions in the ancestor of *A* and *B*. For a given pair of genomes, the estimated 1-cut breakpoint rate is

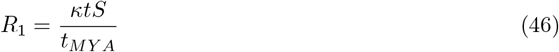

while the 2-cut breakpoint rate is

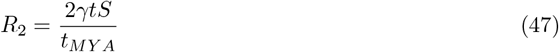

and the hotspot breakpoint rate is

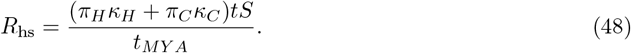

## 5 Acknowledgments

We thank the members of the Nuclear Physics working group in the Center for Theoretical Biological Physics for valuable discussion. Unpublished genome assemblies and sequencing data for all species are used with permission from the DNA Zoo Consortium (dnazoo.org). This research was supported by the National Science Foundation through the awards PHY-2412651 and PHY-2019745, and by the National Institute of General Medical Sciences of the NIH under award R35-GM146852. The content is solely the responsibility of the authors and does not necessarily represent the official views of the funding agencies.

