## Supplemental Information for "Random, fragile, or correlated: Mechanisms of synteny decay in mammals"

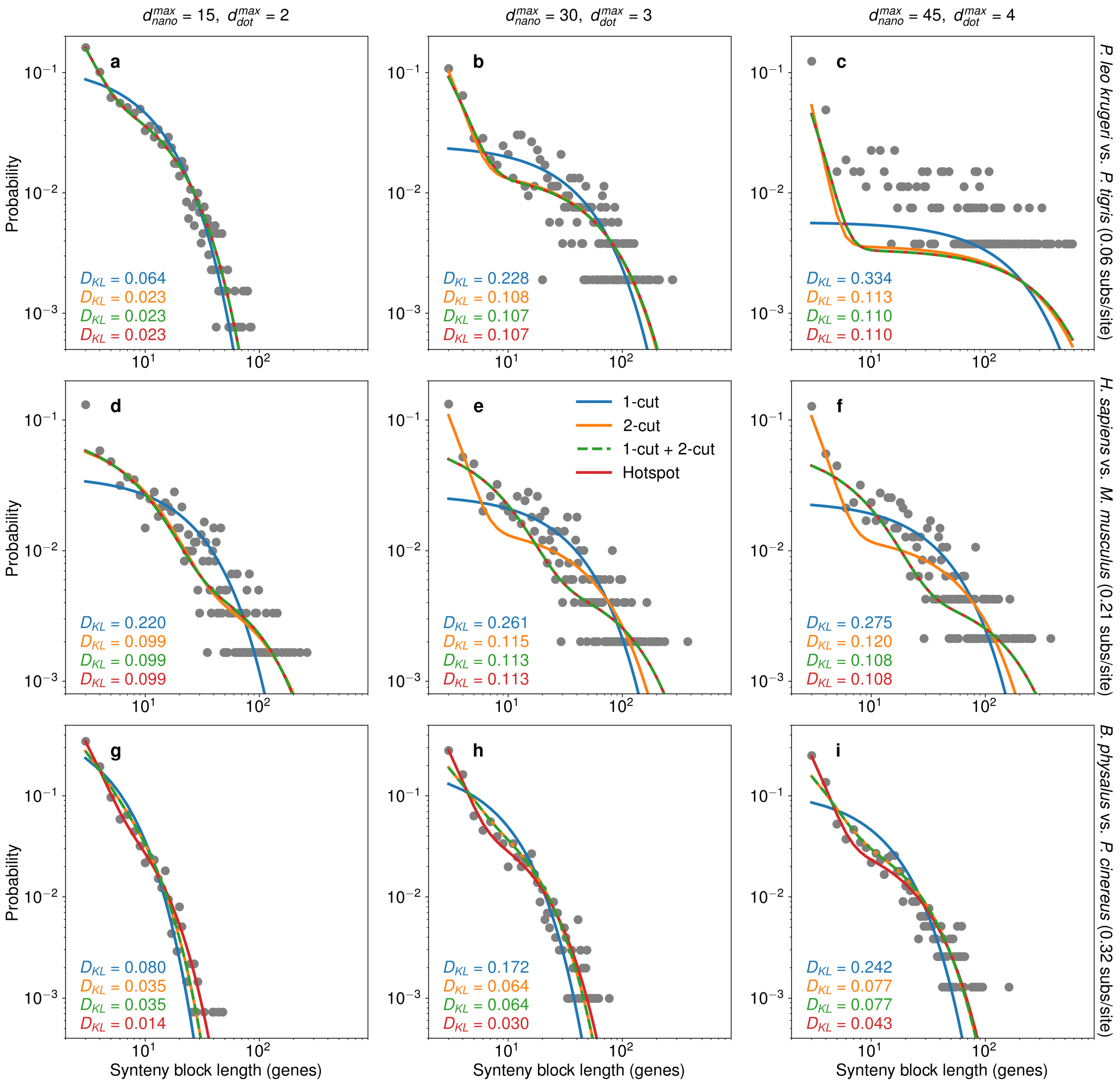


Figure S1: The neutral 1-cut (random breakage model) is inadequate for describing synteny block length distributions across three species pairs and APES parameters. The first column shows empirical distributions and fits for the most conservative parameters tested ($d_{\text{nano}}^{\text{max}} = 15$ and $d_{\text{dot}}^{\text{max}} = 2$), the second column shows the same for intermediate parameters ($d_{\text{nano}}^{\text{max}} = 30$ and $d_{\text{dot}}^{\text{max}} = 3$), and the third column for the most permissive parameters ($d_{\text{nano}}^{\text{max}} = 45$ and $d_{\text{dot}}^{\text{max}} = 4$). The first row shows results for *P. leo krugeri* vs. *P. tigris* (separated by 0.06 amino acid substitutions per site on average), the second row *H. sapiens* vs. *M. musculus* (0.21 amino acid substitutions per site on average), and the third row *B. physalus* vs. *P. cinereus* (0.32 amino acid substitutions per site on average). Color-coded Kullback-Liebler divergences ($D_{KL}$) between each model fit and the empirical distribution are shown in the bottom lefthand corner of each subplot. The hotspot 1-cut model has either the lowest or tied-for-lowest $D_{KL}$ with the empirical distribution in all cases. Note that the visually poor fit in **c** results from low evolutionary distance between the two cats *P. leo krugeri* and *P. tigris*, so that many synteny block sizes are observed only once or not at all and normalization of both the empirical and model distributions leads to a shift between them.

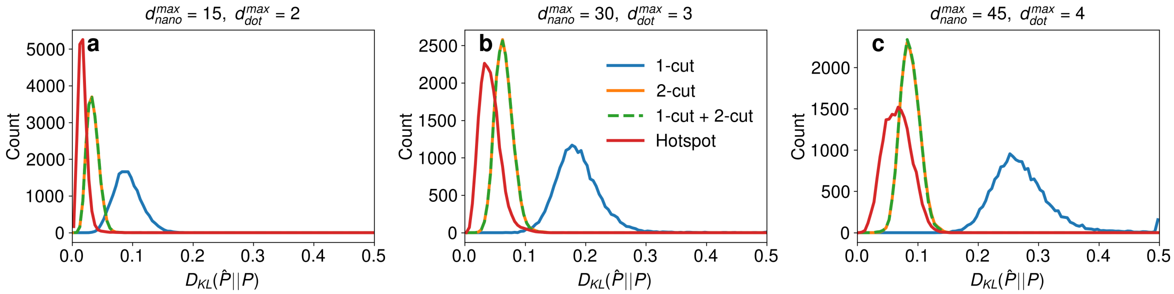


Figure S2: The effects of APES parameters on Kullback-Liebler divergences between actual and theoretical synteny block size distributions ($D_{KL}$). As the APES parameters $d_{\text{nano}}^{\text{max}}$ and $d_{\text{dot}}^{\text{max}}$ increase, the distributions of Kullback-Liebler divergences shift towards larger values, while the relative positions of the four models remain unchanged.

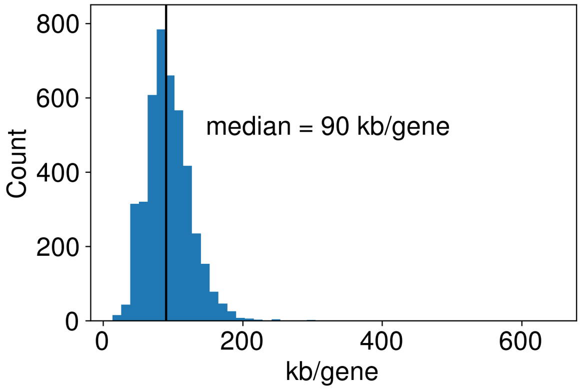


Figure S3: Distribution of kilobases per gene over the 194 genomes in our dataset. For each genome, we simply divided the total length of the genome in kb by the number of genes. For conversion between genes to kb, we use the median value of 90 genes/kb.

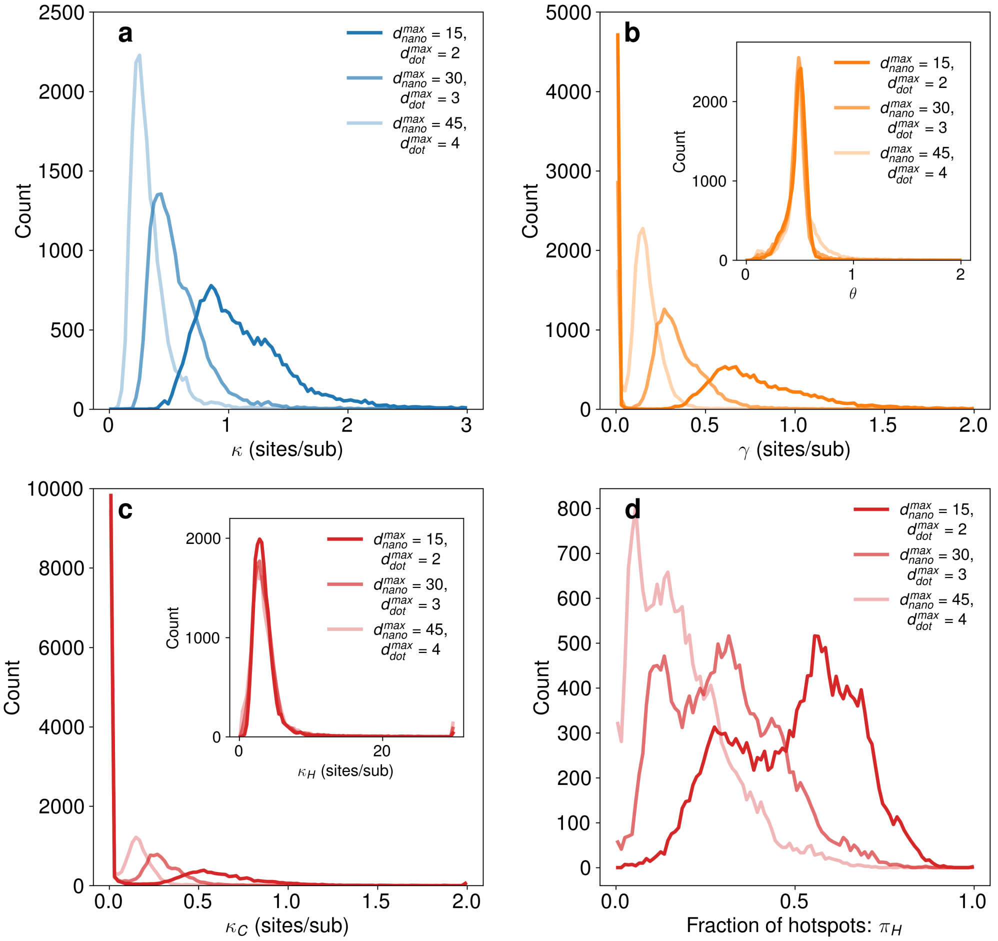


Figure S4: The effects of APES parameters on inferred cut rates and hotspot fractions. For the 1-cut model **a**, the distribution of inferred 1-cut rates ($\kappa$) shifted towards smaller values with increased $d_{\text{nano}}^{\text{max}}$ and $d_{\text{dot}}^{\text{max}}$. This is expected, as larger values of the APES parameters result in fewer, larger synteny blocks with fewer breakpoints. Similarly, the 2-cut rates ($\gamma$) shifted to smaller values with increased APES paramters. Interestingly, the 2-cut size parameter ($\theta$) was largely insensitive to APES parameters (inset). In the hotspot model **c**, the distribution of coldspot cut rates ($\kappa_C$) displayed similar behavior to $\kappa$ and $\gamma$, while the hotspot cut rates ($\kappa_H$) distribution was insensitive to APES parameters. The inferred hotspot fraction of inter-gene regions also shifted leftward with increased APES parameters **d**, with the majority of inter-gene regions being coldspots for $d_{\text{nano}}^{\text{max}} = 30$ and $d_{\text{dot}}^{\text{max}} = 3$ and for $d_{\text{nano}}^{\text{max}} = 45$ and $d_{\text{dot}}^{\text{max}} = 4$.

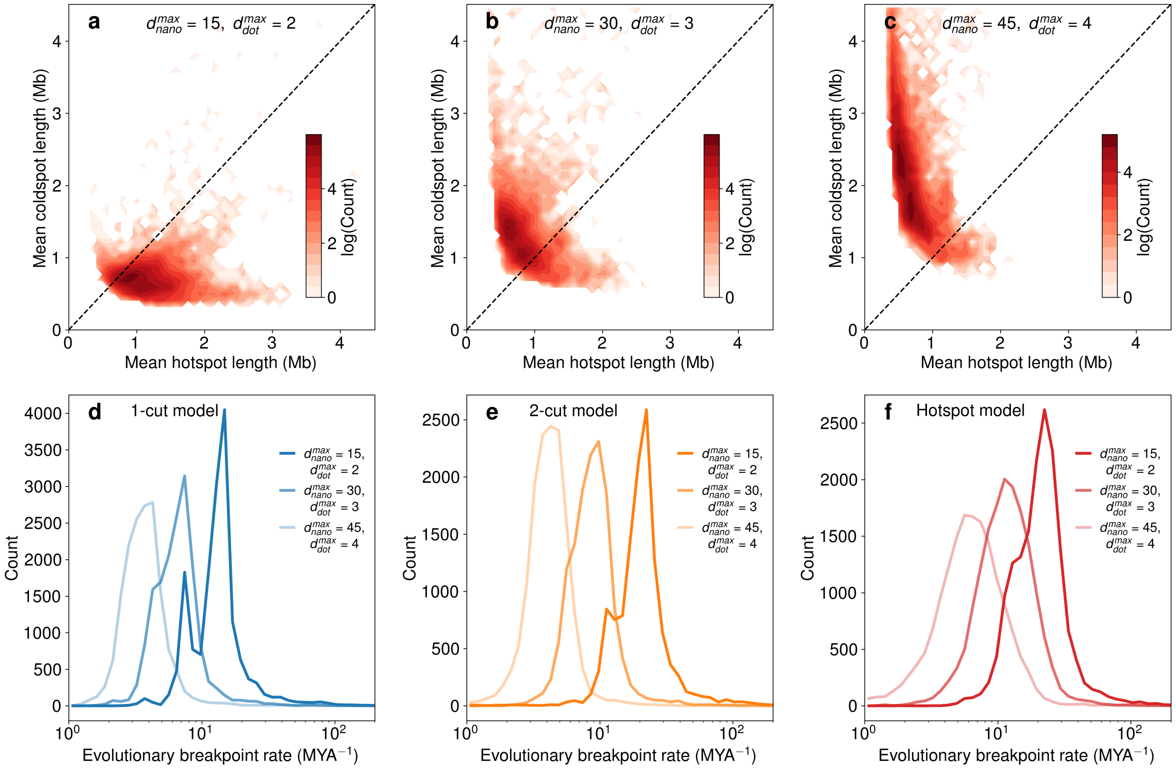


Figure S5: The effects of APES parameters on inferred hotspot and coldspot lengths and breakpoint rates. As APES parameters allow for longer microsynteny blocks from **a** to **b** to **c**, there is a shift from roughly equally-sized hotspots and coldspots to longer coldspots and hotspots smaller than 1 Mb. This result is consistent with the breakpoint rates calculated for the 1-cut model **d**, 2-cut mode **e**, and hotspot model **f** with the same set of APES parameters, where breakpoint rates decrease with larger $d_{\rm nano}^{\rm max}$ and $d_{\rm dot}^{\rm max}$. This is because with larger APES parameters, there are fewer breakpoints and the inferred rates are lower.

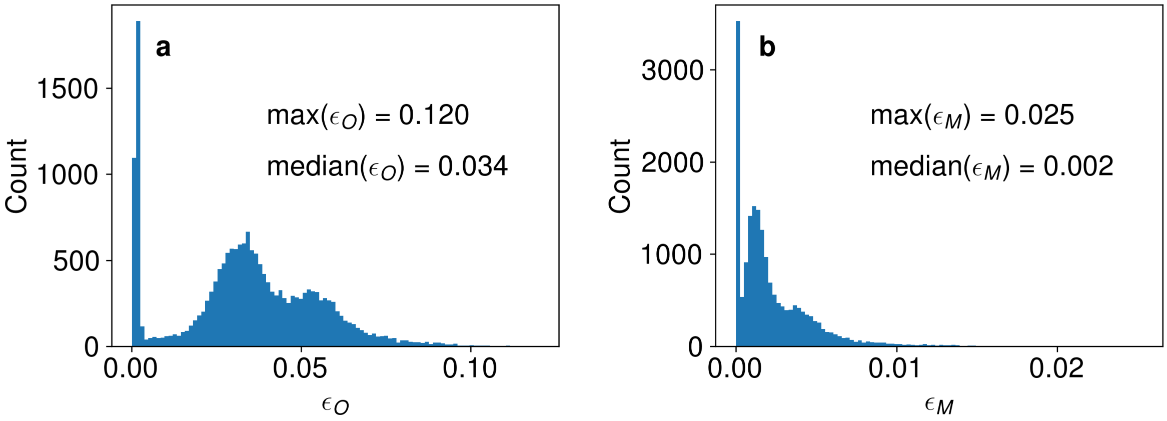


Figure S6: The probabilities of **a** overlapping 2-cuts $\epsilon_O$ and **b** two or more initial 2-cuts at one inter-gene region are low across all fits to pairs of genomes. This indicates that the inferred models are largely internally consistent.
